# Fresh Paraformaldehyde Preserves Thrombus Biochemistry - Infrared Spectroscopy and Secondary Ion Mass Spectrometry Investigation of Pulmonary Artery Thrombi

**DOI:** 10.1101/2025.11.17.688840

**Authors:** Simbarashe Moyo, Andrzej Wróbel, Bartosz Leszczyński, Magdalena Skalska, Jakub Stępniewski, Grzegorz Kopeć, Paweł Moskal, Ewa Stępień

## Abstract

**Background:** Biochemical analyses of pulmonary embolism derived thrombi depend critically on fixative quality.

**Objective:** To quantify the impact of paraformaldehyde (PFA) shelf life on thrombus molecular integrity using attenuated total reflectance Fourier transform infrared (ATR–FTIR) spectroscopy and time-of-flight secondary ion mass spectroscopy (SIMS) combined with principal component analysis.

**Methods:** Ten pulmonary artery thrombi were fixed in either aged PFA (6 months; n = 6) or freshly prepared 4% PFA (n = 4); an in-vitro plasma clot control received the same fresh PFA. Triplicate ATR–FTIR spectra (400–4000 cm^−1^) were collected and vector-normalized to the Amide I band. ToF-SIMS measurements were performed using a 30 keV Bi₃⁺ primary ion beam Fixative chemistry was assessed by the carbonyl absorbance at 1700 cm^−1^.

**Results:** Freshly fixed thrombi displayed narrow Amide I (~1650 cm^−1^) and Amide II (~1540 cm^−1^) bands with a higher signal-to-noise ratio (SNR; median 41.8) than aged-PFA samples (18.2; P < .01). Carbonyl absorbance at 1700 cm^−1^ was markedly higher in fresh PFA. Aged PFA introduces chemical variances of biological samples in low-mass molecule fragments.

PCA showed clear separation of fresh versus aged specimens and alignment of fresh thrombi with plasma controls across spectral windows.

**Conclusions:** PFA solutions older than three months markedly deteriorate thrombus biochemical fidelity. ATR–FTIR and ToF-SIMS offer a rapid quality-control assay prior to molecular analyses.

**Key points:**

I. PFA stored ≥3 months loses >70% of reactive aldehydes, yielding broad Amide envelopes, attenuated lipid bands and a two fold SNR drop.
II. Aged PFA introduces chemical variances of biological samples in low-mass molecule fragments.
III. PCA of fingerprint, protein or lipid windows cleanly separates aged from fresh fixations (PC1 up to 80% variance).
IV. A rapid QC workflow (carbonyl absorbance ≥0.25 a.u. plus PCA verification) safeguards molecular integrity irrespective of patient age, DVT history or clinical severity.
V. Adopting this workflow will harmonise multi centre clot biorepositories and enhance the reproducibility of proteomic, lipidomic and imaging studies.

## 1. Introduction

Paraformaldehyde (PFA) fixation is widely used in preserving biological and medical specimens, namely cellular, and tissue-based molecular imaging workflows; however, fixation-induced chemical alterations remain a critical, underexplored source of spectral variance in high-resolution imaging. Prior work conducted within the Jagiellonian University theranostics and medical physics research frameworks dedicated to the study of the nanostructure of PFA-fixed samples, such as fibrin clots and heart tumors (myxoma) [1–5].

Pulmonary vascular disease is characterized by tightly coupled disturbances in haemostasis, fibrinolysis, inflammation, and endothelial activation. In idiopathic pulmonary arterial hypertension (IPAH), elevated fibrinolytic activity (tPA and plasmin–antiplasmin complexes) alongside increased IL-6 and endothelin-1 shows strong interrelations among these pathways, underscoring the biochemical lability of thrombus-related markers within the pulmonary circulation [6]. At the tissue scale, the architecture of embolic material in acute pulmonary embolism (PE) can inherently limit fibrinolysis. Scanning electron microscopy of a pulmonary artery embolus revealed compact fibrin networks with relatively fewer platelets centrally and erythrocyte-rich peripheral segments, an organization associated with impaired permeability and resistance to lysis [7].

Systemic inflammatory milieus further bias fibrin toward dense, lysis-resistant networks. In chronic obstructive pulmonary disease (COPD), plasma forms clots with reduced permeability, prolonged lysis time, and denser fibrin ultrastructure; these properties correlate with inflammatory burden and are partly amenable to statin therapy [8]. These observations motivate careful biochemical preservation and quality control when analysing pulmonary artery thrombi.

PFA serves as a polymeric formaldehyde source that creates covalent crosslinks stabilising protein structure; with storage, depolymerisation reduces available aldehydes and may compromise preservation [9]. ATR–FTIR spectroscopy provides rapid, label-free biochemical readouts spanning proteins, lipids, and nucleic acids [10–11]. When paired with PCA, small fixation-driven spectral shifts become detectable [12,13]. Given that fixative chemistry can dominate pre-analytical variance, verifying PFA freshness prior to immersion is essential to preserve diagnostically informative features and ensure reproducible molecular phenotyping of PE thrombi. PFA freshness has been studied in brain and tumour biopsies [14], yet its influence on thrombus biochemistry remains poorly characterised. High-resolution time-of-flight secondary ion mass spectrometry (ToF-SIMS) imaging was applied to study molecular cargo patterns observed in nano- and microscale extracellular vesicles to reflect selective biochemical sorting, metabolic responsivity and lipid processing [15–17].

Building on these findings, we hypothesised that aged PFA compromises thrombus molecular preservation, producing spectral signatures discernible by ATR–FTIR and PCA. We extended the ToF-SIMS analysis to directly compare fresh versus chemically aged fixative conditions, to resolve low-mass ion feature contributions driving fixation-state separation in multivariate space and improve the reliability of spectral feature labelling.

## 2. Materials and Methods

### 2.1 Ethics Approval and Consent

The study was approved by the Ethics Committee of Krakowski Szpital Specjalistyczny im. Św. Jana Pawła II (Protocol #2023-152, 14 February 2023). Written informed consent was obtained from each participant or legal representative.

### 2.2 Safety Statement

All protocols involving paraformaldehyde (PFA) were conducted in certified chemical fume hoods, with lab coats, gloves, and safety glasses. Human specimens were handled under Biosafety Level 2 (BSL-2) precautions. All biological material and chemical waste were decontaminated or autoclaved prior to disposal.

### 2.3 Patient Cohort and Thrombus Retrieval

Ten pulmonary artery thrombi were retrieved via catheter-directed mechanical thrombectomy for acute pulmonary embolism between March 2023 and May 2024 at Krakowski Szpital Specjalistyczny im. Św. Jana Pawła II [18–19]. Cohort characteristics are summarized in Table 1, with Full comorbidities in Supplementary Table S1.

**Table 1.**
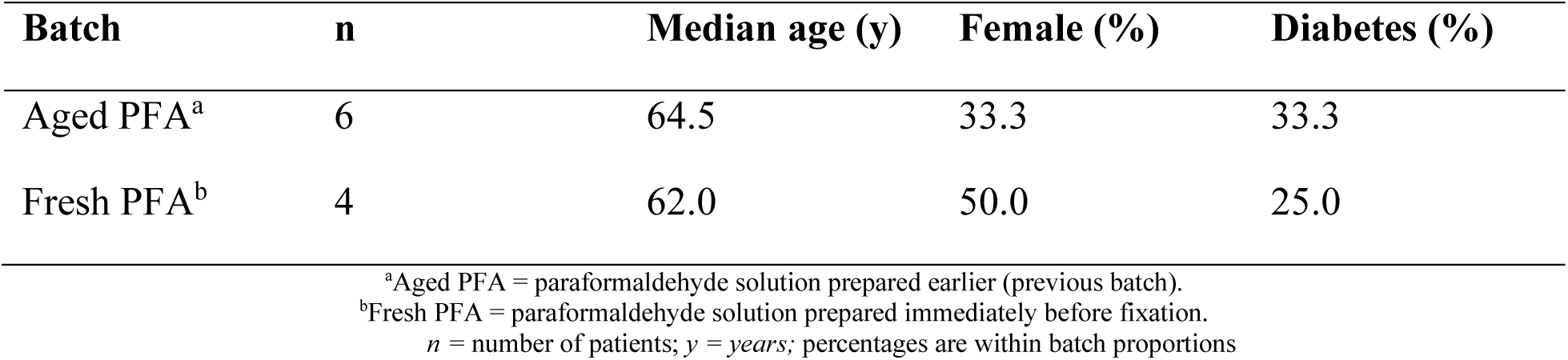
Condensed patient demographics for the two fixation batches.

### 2.4 Baseline Clinical Characteristics and Subgroup Stratification

Table 2 summarizes baseline clinical variables (medians) stratified by history of deep vein thrombosis (DVT; DVT^+^ vs DVT^−^) and by age (≤55 vs >55 years).

**Table 2.** Baseline clinical variables by DVT status and age group (medians; units indicated).

| Variable | DVT Status |  | Age Group |  |
| --- | --- | --- | --- | --- |
|  | DVT<sup>+</sup> (n=5) | DVT<sup>-</sup> (n=5) | ≤55 y (n=4) | >55 y (n=6) |
| sPESI | 2.00 | 2.00 | 2.00 | 2.00 |
| D-dimer baseline (ng/mL) | 10,685 | 4,102 | 9,010 | 6,958 |
| Fibrinogen baseline (g/L) | 3.81 | 3.13 | 3.48 | 3.47 |
| hs-Troponin (ng/L) | 0.04 | 0.04 | 0.04 | 0.06 |
| NT-proBNP (pg/mL) | 5,642 | 2,294 | 5,202 | 3,968 |
| hsCRP (mg/L) | 68.0 | 114.6 | 120.2 | 80.2 |
| Lactate (mmol/L) | 1.90 | 1.20 | 1.80 | 1.60 |

#### Subgroup definitions

Patients were stratified by (i) history of deep vein thrombosis (DVT) thus, DVT^+^ if a prior or concomitant DVT was documented at presentation; DVT^−^ otherwise and (ii) age dichotomized at 55 years. The 55-year cut point was pre specified to improve comparability between groups in this small cohort and to aid interpretability; it was not used for hypothesis testing.

### 2.5 Fixative Preparation and Quality Control

#### Fresh paraformaldehyde (PFA)

Paraformaldehyde powder (4 g) was dissolved in 100 mL 0.01 N phosphate-buffered saline (PBS) at 60 °C with gentle stirring; a few drops of 2 N sodium hydroxide (NaOH) aided dissolution without exceeding 65 °C. Once clear, the pH was adjusted to 7.2–7.4 with 1 M hydrochloric acid (HCl). The solution was sterile-filtered (0.22 μm) and stored at 4 °C (≤30 days) or −20 °C (≤90 days). **Aged PFA**: An identical solution stored at room temperature for six months. **QC metrics:** Each batch followed the standard operating procedure in Table 3; carbonyl absorbance at 1700 cm^−1^ was measured by ATR-FTIR.

**Table 3.** Fixative quality control standard operating procedure.

| Step | Measurement | Acceptance Criterion | Rationale |
| --- | --- | --- | --- |
| 1 | Visual inspection | Clear, colorless | Initial degradation screen |
| 2 | ATR–FTIR carbonyl peak | Absorbance $\geq 0.25$ | Free aldehyde content |
| 3 | pH at 25°C | 7.2–7.4 | Optimal cross-linking |
| 4 | Shelf-life | 90 days | Empirical integrity limit |

### 2.6 Sample Fixation and Storage

Each thrombus was fixed for 24 h at 4 °C in either aged PFA (Batch I, n=6) or fresh PFA (Batch II, n=4), washed (3 × 10 min PBS), and stored at 4 °C for up to three months. Chemical composition and spectra for all thrombi and PFA solutions are presented in Fig. 1, with anonymized labels corresponding to each specimen.

**Figure 1.**
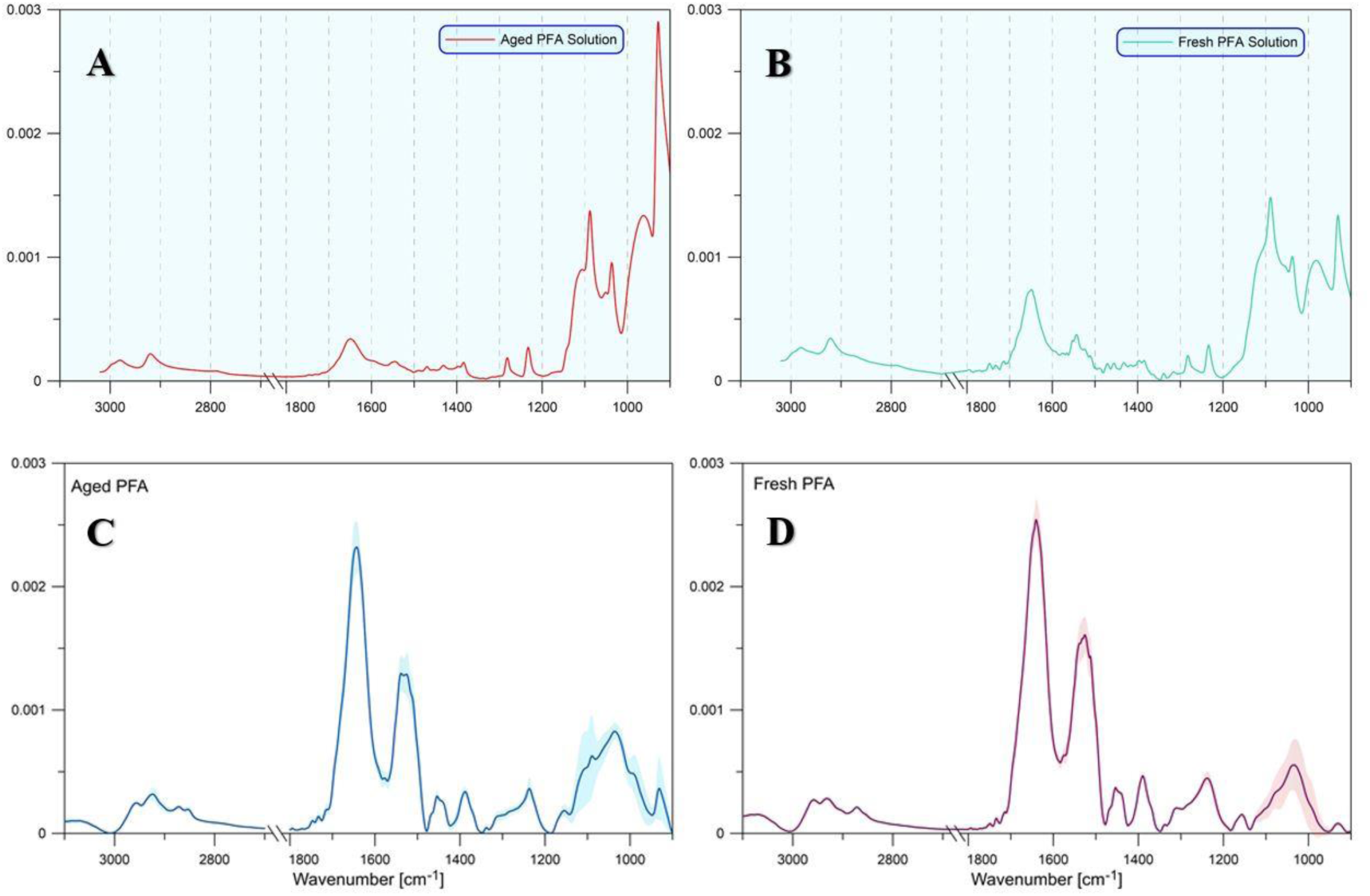
ATR–FTIR of PFA solutions and averaged spectral profiles. **(A)** Aged PFA. The carbonyl band near 1700 cm^−1^ is diminished and broadened with baseline drift, indicating aldehyde depletion. **(B)** Fresh PFA. The carbonyl band is strong and well defined on a flat baseline, indicating preserved chemistry. **(C)** Aged PFA (mean ± SD). Across 1800–1650 cm^−1^, the carbonyl absorbance is lower with greater variability; modest fingerprint features appear at 1200–1000 cm^−1^. **(D)** Fresh PFA (mean ± SD). The carbonyl is higher and sharper with tighter variability; only minor fingerprint features are present. Overall, fresh PFA retains reactive carbonyls, whereas aged PFA shows characteristic degradation signatures.

### 2.7 Plasma Clot Control

Platelet-poor plasma from the Regional Centre for Blood Donation and Blood Treatment in Kraków was thawed and mixed with a thrombin working solution in Tris-HCl buffer. After incubation at 37 °C for 30–60 min, clots were washed, fixed in freshly prepared 4% PFA for at least 2 hours at room temperature or overnight at 4 °C, then rinsed three times (10 minutes each) with Tris-HCl buffer. Dehydration was performed in a vacuum desiccator (relative humidity ≈ 2%) over anhydrous calcium sulfate. Mass was monitored until stable (~4 days) prior to ATR–FTIR analysis.

### 2.8 ATR–FTIR Spectroscopy

Dried samples (~10 mg) were ground, placed on a diamond ATR crystal (Nicolet 6700), and scanned (400–4000 cm^−1^, 128 scans, 4 cm^−1^ resolution). Spectra were averaged, atmospheric-compensated, rubber-band corrected, smoothed (Savitzky–Golay, 2nd order, 9-point window), and vector-normalized to the Amide I band.

#### Deep vein Thrombosis subgroup

Thrombi were classified as originating from patients with a documented history of lower-limb DVT (were additionally grouped by history of deep-vein thrombosis (DVT) and by age (≤55 vs >55 years) for averaged spectra using OriginLab 2025.

### 2.9 Materials and Methods

ToF-SIMS measurements were performed using a 30 keV Bi₃⁺ primary ion beam under ultra-high vacuum on the analytical platform to profile molecular fragments from biological specimens fixed under fresh and chemically aged paraformaldehyde conditions. In static mode, the surface integrity was preserved by maintaining an ion fluence of ≤ 1 × 10^12^ ions/cm^2^, while additional fast-imaging acquisitions enabled high-throughput ion-distribution reconstruction. Each sample was analysed in both positive and negative ion modes (4 = 3 replicates per mode; 250 × 250 µm^2^ ROI, 128 × 128 pixels), with internal calibration applied using reference carbon-cluster and hydride ions. Spectral acquisitions up to m/z 911 amu were normalized to total ion counts prior to statistical analysis.

### 2.9 Principal Component Analysis

PCA was conducted on three spectral regions: the biochemical fingerprint window (900–1800 cm^−1^), the protein-associated Amide I+II region (1480–1800 cm^−1^), and the lipid-rich C–H stretch window (2750–3010 cm^−1^). All spectra were vector-normalized to Amide I prior to analysis. Components with eigenvalues ≥ 1 (Kaiser criterion) were retained. Score plots (PC1 vs PC2) were shown with a 95% Hotelling’s T^2^ ellipse.

### 2.10 Statistical Analyses

Analyses were performed in Python (SciPy 1.13). For each sample, spectral quality was quantified as SNR, computed as the Amide I peak height (~1650 cm^−1^) divided by the RMS of baseline noise estimated over 1900–2000 cm^−1^ after preprocessing. SNR is summarized as median [IQR]. Because SNR and clinical variables exhibited non-Gaussian features, between-batch differences were assessed using two-tailed Mann–Whitney U tests. Associations between SNR and clinical markers were evaluated with Spearman’s ρ using exact P values and 95% bootstrap confidence intervals. Mass spectrometry data were evaluated using Shapiro–Wilk, Levene, ANOVA, FDR-corrected Tukey, and database-supported molecular annotation pipelines, according to mass-spectrometry imaging and lipidomic reporting standards. All tests were two-sided with α = .05; no imputation was applied, and ties used SciPy defaults.

## 3 Results

### 3.1 PFA Quality Analysis

#### 3.1.1 Fixative Chemical Stability

Fresh and aged PFA solutions exhibited markedly different carbonyl signatures. Fresh PFA produced a strong absorbance at 1700 cm^−1^ (~0.31 a.u.), reflecting abundant free aldehyde groups essential for cross-linking. In contrast, aged PFA displayed a greatly diminished carbonyl peak (~0.09 a.u.), indicating >70% loss of reactive species after prolonged storage. Additional broadening and baseline drift between 1650 and 1800 cm^−1^ suggest polymerization and oxidation.

#### 3.1.2 Averaged Spectral Profiles of Fixed Thrombi

To assess whether PFA quality alters thrombus chemistry, we applied the same ATR–FTIR analysis to thrombi fixed in aged versus fresh PFA. Quantitative analysis of averaged spectra revealed systematic differences between fixation conditions across major biochemical regions (Fig. 1C, D), demonstrating the molecular consequences of PFA degradation on preservation quality.

##### Aged-PFA Group (n=6)

Thrombi fixed in degraded PFA (specimens 1EDU, 1DKU, 1EA, 1FDA, 1CFL, 1BM1) showed compromised protein preservation with broad Amide I peaks centered at 1649 ± 3 cm^−1^. The broadening reflects disrupted hydrogen bonding networks and alters secondary structure due to inadequate cross-linking [14,20].

##### Fresh-PFA Group (n=4)

Thrombi preserved in fresh PFA (specimens 1GIN, 1PAN, 1ZAN, 1ZMA) exhibited sharp Amide I peaks at 1652 ± 1 cm^−1^ and well-defined Amide II bands at 1540 ± 2 cm^−1^, consistent with preserved protein conformation and effective aldehyde mediated cross-linking.

#### 3.1.3 Chemometric Discrimination by PCA (Fixative freshness)

The composite PCA figure (Fig.2) summarizes three windows. **(A)** Fingerprint (900–1800 cm^−1^): PC1 69.0% (PC2 14.7%) separates aged-PFA (negative PC1) from fresh-PFA and plasma controls (positive PC1). **(B)** Amide I+II (1480–1800 cm^−1^): discrimination sharpens (PC1 78.0%, PC2 11.5%), indicating predominant protein-band perturbation. **(C)** Lipid C–H stretch (2750–3010 cm^−1^): separation persists (PC1 80.5%, PC2 15.6%), consistent with CH₂/CH₃ degradation when aldehydes are depleted. Plasma controls align with fresh-PFA.

**Figure 2.**
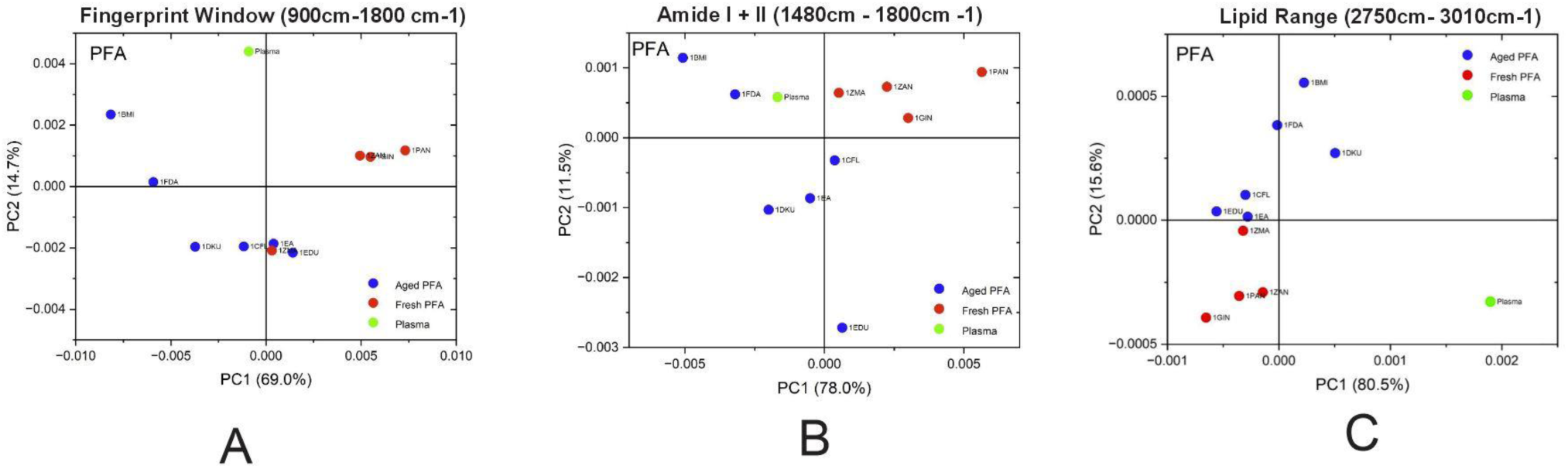
Principal component analysis (PCA) of attenuated total reflectance Fourier transform infrared (ATR-FTIR) spectra. **(A)** Fingerprint region (900–1800 cm^−1^). Principal component 1 (PC1) explains 69.0% of the variance and principal component 2 (PC2) explains 14.7%; aged paraformaldehyde (PFA; blue) separates from fresh PFA (red) and plasma (green). **(B)** Amide I and II region (1480–1800 cm^−1^). PC1 explains 78.0% and PC2 11.5%; the increased discrimination highlights protein sensitivity to fixative freshness. **(C)** Lipid C–H stretch region (2750–3010 cm^−1^). PC1 explains 80.5% and PC2 15.6%; separation persists, driven by CH₂/CH₃ stretching contributions.

#### 3.1.4 Signal-to-Noise Analysis

Fresh-PFA thrombi showed a median SNR of 41.8 vs 18.2 for aged-PFA thrombi (P<0.01), a more than two-fold difference exceeding inter-patient variability.

### 3.2 Spectral Comparison by DVT History

Across 900–3100 cm^−1^, DVT^+^ (n = 5) and DVT^−^ (n = 5) traces were virtually superimposable. No Amide I/II shifts or lipid to protein ratio changes, phosphate at 1080 cm^−1^ was unchanged. Median SNR did not differ (40.9 vs 40.6; P = .10).

**Figure 3.**
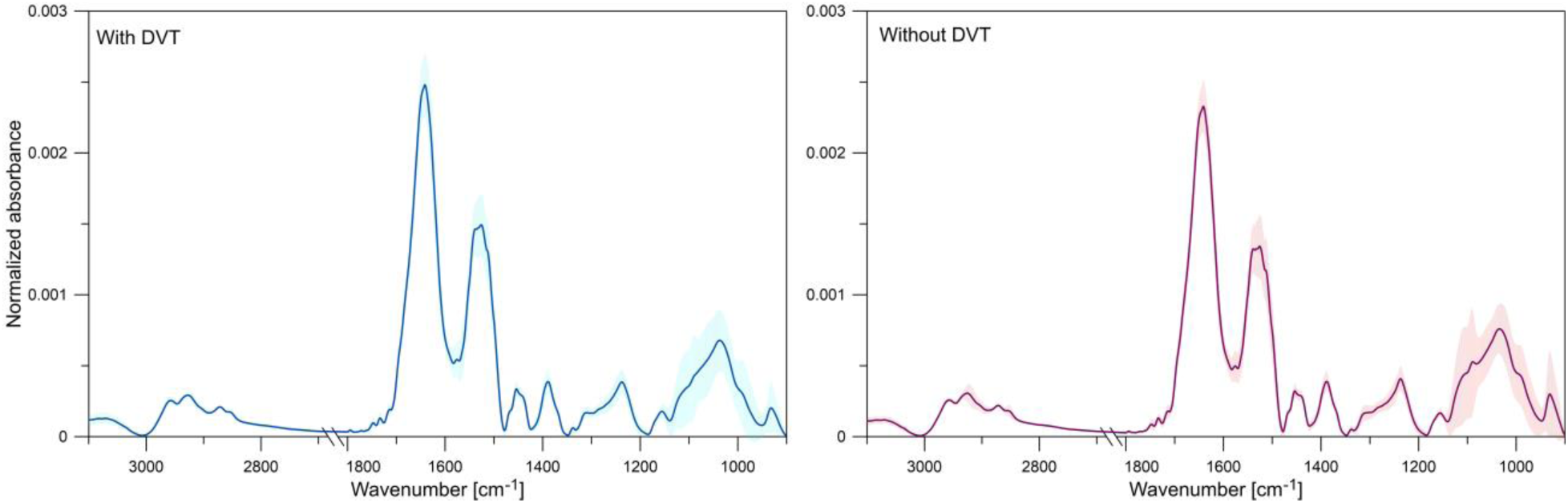
Averaged ATR–FTIR spectra grouped by history of deep-vein thrombosis (DVT). Black trace = DVT^+^ thrombi (n=5); cyan = DVT^−^ thrombi (n=5). Shaded bands denote ± 1 SD.

### 3.3 Spectral Comparison by Age Group

In Fig 4, Younger (≤55 y) and older (>55 y) cohorts showed near complete spectral overlap with indistinguishable SNR (40.3 vs 40.2; P = .97); no age-related clustering was seen in PCA. This aligns with reports that age-related lipid metabolism shifts are subtle and context-dependent [20,21].

**Figure 4.**
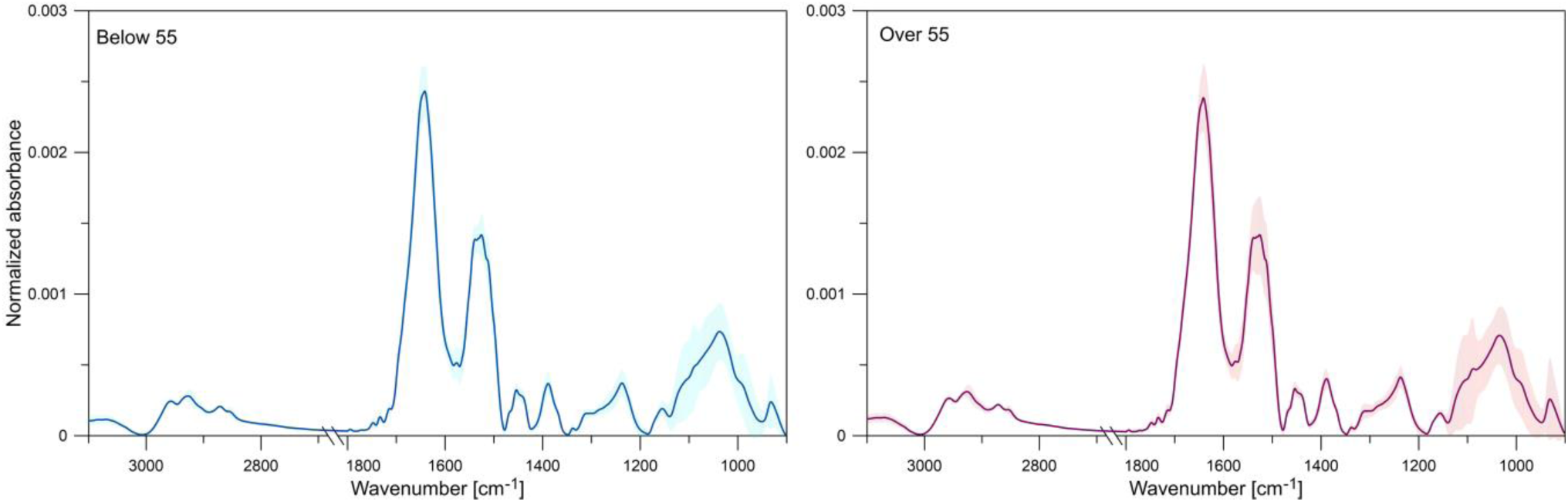
Averaged ATR–FTIR spectra grouped by age. Cyan trace = patients ≤ 55 y (n=4); magenta = patients > 55 y (n=6). Shaded bands denote ± 1 SD.

#### 3.3.1 Chemometric Discrimination by PCA (DVT vs Age, composite)

The composite PCA summary (Fig. 5) contrasts deep vein thrombosis (DVT) history and age across three spectral windows using identically pre-processed, Amide I-normalised spectra. In every window, scores from the clinical subgroups are extensively intermixed with similar dispersion, indicating that subgroup membership does not explain the sample variance once fixative freshness is controlled.

**Figure 5.**
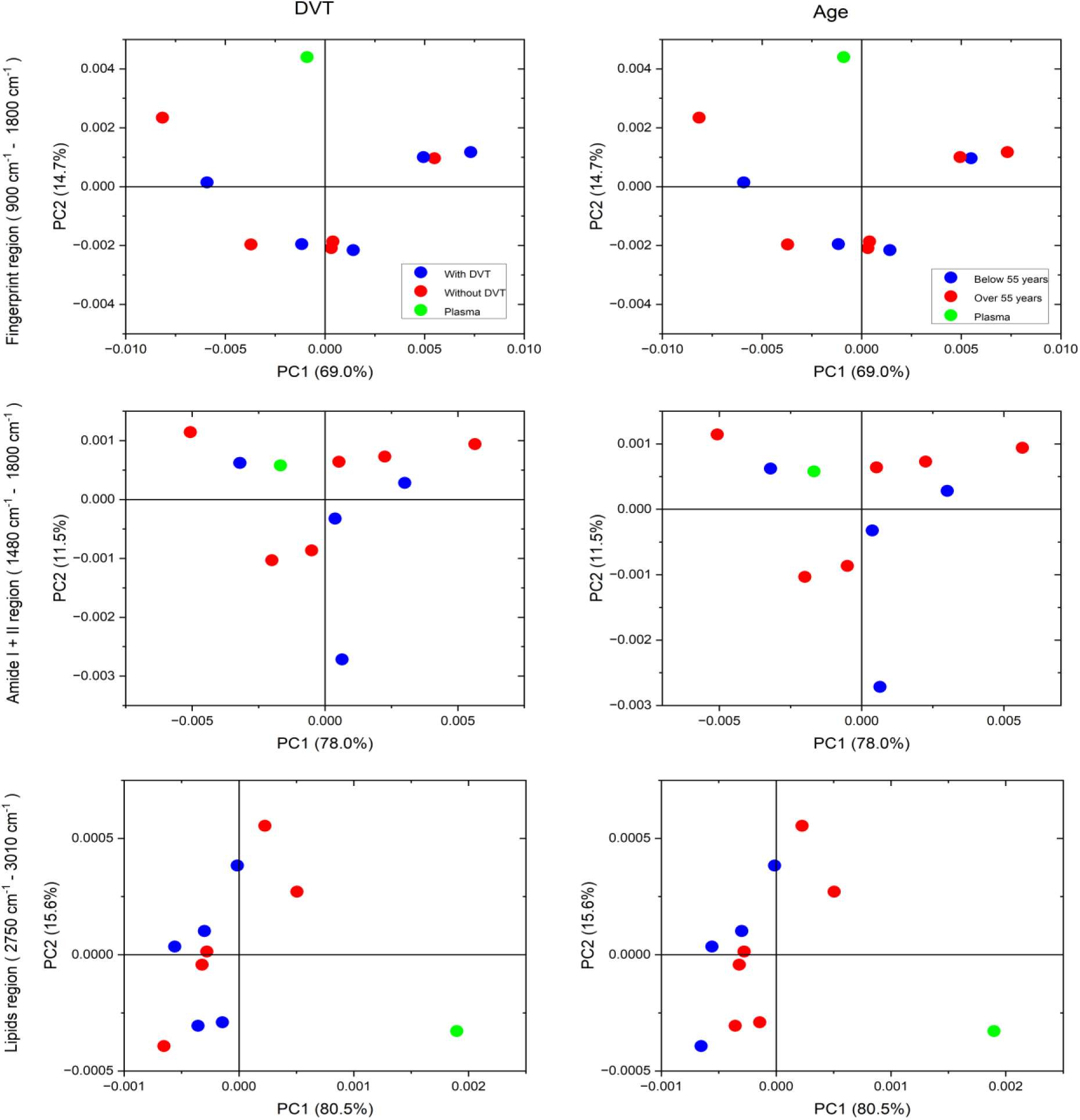
Composite principal component analysis (PCA) comparing DVT history and age across spectral windows. *Left column (DVT comparison):* Blue = DVT⁺ (n=5); Red = DVT⁻ (n=5); Green = plasma control. *Right column (age comparison):* Blue = ≤55 years (n=4); Red = >55 years (n=6); Green = plasma control. Rows correspond to the fingerprint region (900–1800 cm^−1^), the Amide I+II region (1480–1800 cm^−1^), and the lipid C–H stretch region (2750–3010 cm^−1^). The percentages of variance explained by PC1 and PC2 are shown on the axes; scores largely overlap across groups.

##### DVT comparison (left column)

In the fingerprint region (900–1800 cm^−1^; PC1 = 69.0 %, PC2 = 14.7 %), DVT⁺ (n = 5) and DVT⁻ (n = 5) occupy overlapping areas with no axis-wise trend. Restricting to the protein bands (Amide I+II, 1480–1800 cm^−1^; PC1 = 78.0 %, PC2 = 11.5 %) tightens the cloud yet maintains full overlap. In the lipid C–H stretch window (2750–3010 cm^−1^; PC1 = 80.5 %, PC2 = 15.6 %), patient samples cluster near the origin without DVT-related separation, while plasma controls tend to lie at positive PC1.

##### Age comparison (right column)

Age-stratified scores (≤ 55 y, n = 4 vs > 55 y, n = 6) similarly intermix across the fingerprint, protein, and lipid windows, with comparable within-group dispersion and no monotonic gradient along PC1 or PC2.

##### Overall

Across windows, neither prior DVT nor age yields separable clusters when fixative freshness is standardised, consistent with the axis-wise variance shown on each plot.

### 3.5 High-resolution secondary mass spectrometry characrtisation of chemical modifications

The total ion counts normalized high-resolution ToF-SIMS spectra revealed distinct low-mass ion feature contributions associated with the fixation state (**Figure 6**). PCA score plots demonstrated reproducible separation between the aged (A₁–A₅) and fresh (F₁–F₅) conditions in both ionization modes, confirming fixation-dependent spectral shifts. PC1 loadings limited to m/z ≤ 200 amu indicated that Na⁺, C-N/O-related fragments, and phosphate-containing clusters were the primary drivers of multivariate clustering. Paired boxplots confirmed directional intensity shifts for matched m/z signals between groups, supporting downstream statistical comparability without sample-level identification. All 2D ion distribution maps obtained in the negative ion mode are provided in supplementary material (Fig. S1), and all corresponding maps from the positive ion mode are included to support full spatial-throughput visualization beyond the main results panel (Fig. S2).

**Figure 6.**
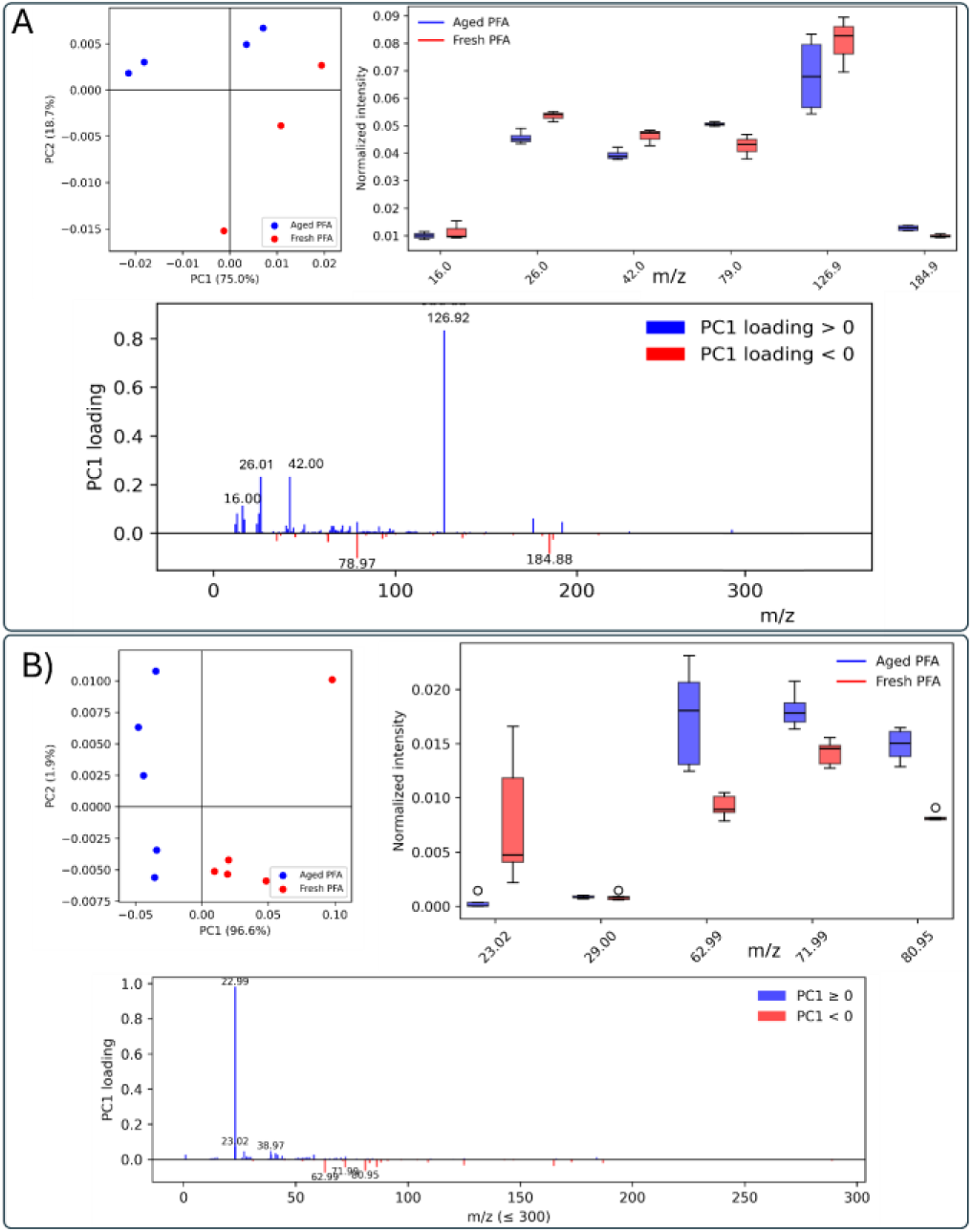
Overview of ToF-SIMS results for biological specimens fixed under fresh (F) and chemically aged (A) paraformaldehyde conditions. **A) negative** ion mode and **B) positive** ion mode: (i) score plot of PC1 vs. PC2 calculated from normalized spectra of samples fixed with aged PFA (blue circles) and freshly prepared PFA (red circles)showing group clustering, (ii) PC1 loading plots limited to the informative low-mass range (≤ 200 amu) with key *m/z* peak labels, and (iii) paired boxplots displaying TIC-normalized ion intensity distributions for matched Aged vs. Fresh groups for each labeled *m/z*. All data reflect measurements acquired under identical analytical conditions on a 250 × 250 µm^2^ ROI, all spectra were normalized to total ion intensity prior to multivariate analysis, and acquisition parameters (primary ion species, dose, pixel size and field of view) were kept identical across fixation conditions.

### 3.4 Clinical Parameter Correlation with Spectral Quality

Spearman analysis showed no meaningful association between SNR and routinely collected markers including simplified Pulmonary Embolism Severity Index (sPESI), baseline D-dimer, fibrinogen, high-sensitivity troponin (hs-troponin), N-terminal pro–B-type natriuretic peptide (NT-proBNP), lactate, high-sensitivity C-reactive protein (hsCRP), or neutrophil-to-lymphocyte ratio (NLR) (all |ρ| < 0.25, P > 0.20). Median IQR values by fixative batch appear in Table 1. Fixative freshness remained the sole significant driver of SNR (fresh PFA: 41.8 vs aged PFA: 18.2; P < 0.01).

**Table 4.** Key clinical variables and attenuated total reflectance Fourier transform infrared (ATR-FTIR) signal-to-noise ratio by fixative batch. Data are median [IQR].

| Variable | Fresh paraformaldehyde (PFA; n=4) | Aged PFA (n=6) |
| --- | --- | --- |
| sPESI | 2.0 | 2.0 |
| D-dimer baseline (ng/mL) | 10,712 | 5,691 |
| Fibrinogen baseline (g/L) | 3.59 | 3.48 |
| hs-Troponin (ng/L) | 37.5 | 172 |
| NT-proBNP (pg/mL) | 2,284 | 7,570 |
| hsCRP (mg/L) | 110.5 | 91.3 |
| NLR | 4.51 | 5.43 |
| Lactate (mmol/L) | 1.60 | 1.80 |
| **SNR** | **41.8** | **18.2** |

**Table 5.** Signal-to-noise ratio (SNR) by deep vein thrombosis (DVT) status and by age group. Medians [IQR]; P values from two-tailed Mann–Whitney U tests.

DVT Status
| **Group** | **n** | **SNR** | **P value** |
| --- | --- | --- | --- |
| DVT<sup>+</sup> | 5 | 40.9 | 0.10 |
| DVT<sup>−</sup> | 5 | 40.6 |  |

Age Group
| **Group** | **n** | **SNR** | **P value** |
| --- | --- | --- | --- |
| ≤55 y | 4 | 40.3 | 0.97 |
| >55 y | 6 | 40.2 |  |

Median SNR did not differ by DVT history (40.9 vs 40.6; P = 0.10) or age (55y vs >55y: 40.3 vs 40.2; P = 0.97). Consistently, composite PCA (Fig. 5) shows extensive overlap across all windows, plasma controls cluster tightly. Thus, once fixative freshness is standardized, neither prior DVT nor age produces detectable biochemical shifts in thrombus spectra.

## 4. Discussion

### 4.1 Principal Findings

Across fingerprint (900cm−1 to 1800cm−1), protein (Amide I+II; 1480cm−1 to 1800cm−1) and lipid (CH-stretch; 2750cm−1 to 3010cm−1) windows, fixative freshness specifically, preservation of reactive aldehyde groups was the dominant determinant of thrombus ATR–FTIR integrity. Aged 4% PFA (>3 months) generated broad Amide envelopes, attenuated CH2/CH3 bands and a two fold SNR loss, whereas freshly prepared PFA conserved sharp protein and lipid-associated peaks. PCA verified this chemistry driven separation: PC1 explained 69% (fingerprint), 78% (protein) and 80% (lipid) of total variance.

Consistent with these observations, the aged fixative failed the pre specified quality control criteria its carbonyl absorbance at 1700cm−1 fell below 0.25a.u. and the solution exceeded the 90 day shelf life confirming its unsuitability for reliable biochemical preservation.

The ToF-SIMS spectral variability observed between fixation conditions confirms that chemically aged and fresh PFA states introduce structured, low-mass fragment differences detectable in multivariate space. Restricting PC1 loadings to ≤ 200 amu showed on Figure 1 provides a high signal-to-noise window for publication-grade ion feature labeling, reducing visual distortion while preserving separation drivers. The paired boxplot strategy further contextualised PC1-defining ions, enabling direct Aged vs Fresh intensity comparability consistent with MSI reporting conventions. This panel supports the manuscript’s narrative that fixation state shapes spectral clustering without confounding sample identifiers in main PCA projections.

Altogether, this refined low-mass spectral window provides a robust basis for ion-map reconstruction and fixation-aware feature reporting in nanoscale tissue characterisation.

### 4.2 Impact on Specific Molecular Classes

Aged PFA, depleted of >70% of free aldehydes, cross-links inefficiently; thrombi exhibit broadened Amide I/II envelopes and baseline drift, consistent with altered secondary structure [14,23]. The lipid window showed the greatest PC1 variance (80.5%), reflecting erythrocyte membrane and platelet-derived lipid content. Degraded fixative (via methanol/formic acid) can extract lipids, reducing CH2/CH3 intensity; this is concordant with lipidomics links to thrombus biology [24–27]. Mixed contributions. Intermediate sensitivity (PC1 ≈69%) arises from concurrent perturbations to protein, lipid, and carbohydrate bands.

### 4.3 Clinical Implications

A 1min attenuated total reflectance–Fourier transform infrared (ATR–FTIR) scan of the fixative, combined with a carbonyl absorbance threshold (≥0.25 arbitrary units [a.u.]), effectively identifies degraded paraformaldehyde (PFA) before immersion, thereby preventing artefacts in protein, lipid, and mixed bands. Given that spectra are unaffected by age, deep vein thrombosis (DVT) history, or clinical severity, a chemistry-centered quality control (QC) protocol standardizes sample quality across centers without the need for demographic stratification. Fresh PFA maintains labile fibrin structures and membrane lipids, thereby minimizing proteolytic and extraction artefacts that could complicate quantitative mass spectrometry and imaging workflows.

### 4.4 Limitations

Sample size was modest (ten thrombi; six aged-PFA, four fresh) and evaluated a single PFA concentration (4%). Only one fixation temperature and a single ATR platform were used.

### 4.5 Comparison with Previous Work

Fixation-related artefacts like those observed here have been reported in brain and tumour tissues [12,14]. To our knowledge, this study provides the first systematic evaluation of such artefacts in pulmonary artery thrombi from patients with pulmonary embolism (PE). Our observations align with the established understanding that dense fibrin architectures impede fibrinolysis [1–3] and confirm usefulness of vibration spectroscopy for thrombi analysis [28].

### 4.6 Future Directions

Building on these results, ongoing work will (i) assess microwave-enhanced fixation to shorten immersion times, (ii) test alternative fixatives such as glyoxal and zinc formalin, and (iii) integrate partial least squares discriminant analysis (PLS-DA) to predict fixative age directly from raw attenuated total reflectance–Fourier transform infrared (ATR–FTIR) spectra. Together, these efforts aim to refine fixation protocols, minimize spectral distortions, and broaden the applicability of vibrational spectroscopy for accurate biochemical characterization of thrombi. Supplementing these studies with complementary methods using positronium lifetime spectroscopy and secondary ion mass spectrometry (SIMS) will allow for the assessment of the nanostructure of the fibrin clot [5,16,17].

## 5 Conclusions

Fixative chemistry, not clinical status, dominates pre-analytical variance in molecular thrombus profiling. A QC workflow combining fixative carbonyl assessment with ATR–FTIR PCA reflects clinically informative protein and lipid features and -enables reliable, multi centre discovery science.

## Ethics, Transparency, and Reproducibility

### IRB Approval

Krakowski Szpital Specjalistyczny im. Św. Jana Pawła II, Protocol #2023-152.

### Informed Consent

Obtained from all participants or legal representatives.

### Conflict of Interest

The authors declare no competing interests.

### Funding

Foundation for Polish Science (TEAM POIR.04.04.00-00-4204/17); National Science Centre of Poland (2021/42/A/ST2/00423; 2025/57/N/NZ7/02297); Jagiellonian University Excellence Initiative.

### AI-Usage Disclosure

AI was used under human supervision solely for language refinement; scientific content verified by the authors.

## CRediT Author Statement

S. Moyo — Investigation; Writing – Original Draft; Methodology

A. Wróbel — Formal Analysis; Visualization; Investigation; Methodology

B. Leszczyński — Investigation; Methodology

M. Skalska — Investigation; Methodology; Writing – Review & Editing

J. Stępniewski — Clinical Investigation; Patient Recruitment; Resources

G. Kopeć — Clinical Investigation; Patient Recruitment; Resources

P. Moskal — Supervision

E. Stępień — Conceptualization; Writing – Review & Editing; Funding Acquisition; Supervision

## Notes

### Competing Interest Statement

The authors have declared no competing interest.

### Summary of Updates

This revised version of the manuscript incorporates new high-resolution ToF-SIMS analyses to strengthen the characterization of fixation-dependent chemical changes in pulmonary artery thrombi. A dedicated subsection has been added describing total ion count normalized spectra, PCA score and loading plots, and paired boxplots comparing fresh versus chemically aged paraformaldehyde fixation conditions. These additions clarify how low-mass ion features, including Na^+, CN/CO fragments, and phosphate-containing clusters, drive multivariate separation between fixation states. Figure 6 has been fully updated to present the expanded ToF-SIMS results in both ion modes, with consistent acquisition and normalization details provided for reproducibility. Supplementary materials now include comprehensive 2D ion distribution maps for negative and positive ion modes. Methods have been refined to provide clearer descriptions of fixation protocols, spectroscopy acquisition parameters, and preprocessing steps for ToF-SIMS, ATR-FTIR, and PCA. Minor edits were made to the Abstract to reflect the new findings, and text throughout the Results and Discussion has been clarified for improved coherence and interpretation regarding biochemical preservation under fresh PFA fixation. References and supplemental files have been updated, and minor editorial corrections applied throughout. These revisions collectively enhance the analytical depth, transparency, and overall clarity of the manuscript while maintaining its scientific objectives.

## References

1. Stepien EL, Kwaśniewska M, Rębowska E, Golański J, Drygas W. Modified thrombin formation and fibrinolysis in an ultra-endurance marathon swimmer. Scand J Med Sci Sports. 2017 May;27(5):567–570. doi: 10.1111/sms.12836.

2. Stępień E, Kabłak-Ziembicka A, Musiałek P, Tylko G, Przewłocki T. Fibrinogen and carotid intima media thickness determine fibrin density in different atherosclerosis extents. Int J Cardiol. 2012 Jun 14;157(3):411–3. doi: 10.1016/j.ijcard.2012.03.140.

3. Stępień E, Miszalski-Jamka T, Kapusta P, Tylko G, Pasowicz M. Beneficial effect of cigarette smoking cessation on fibrin clot properties. J Thromb Thrombolysis. 2011 Aug;32(2):177–82. doi: 10.1007/s11239-011-0593-6.

4. Moskal P, Kubicz E, Grudzień G, Czerwiński E, Dulski K, Leszczyński B, Niedźwiecki S, Stępień EŁ. Developing a novel positronium biomarker for cardiac myxoma imaging. EJNMMI Phys. 2023 Mar 24;10(1):22. doi: 10.1186/s40658-023-00543-w.

5. Moyo S, Moskal P, Stępień E. Feasibility study of positronium application for blood clots structural characteristics. Bio-Algorithms and Med-Systems 2022; 18(1): 163–167. doi: 10.2478/bioal-2022-0087.

6. Kopeć G, Moertl D, Steiner S, Stępień E, Mikołajczyk T, Waligóra M, et al. Markers of thrombogenesis and fibrinolysis and their relation to inflammation and endothelial activation in patients with idiopathic pulmonary arterial hypertension. PLoS One. 2013; 8(12): e82628. doi: 10.1371/journal.pone.0082628.

7. Undas A, Stępień E, Rudziński P, Sadowski J. Architecture of a pulmonary thrombus removed during embolectomy in a patient with acute pulmonary embolism. J Thorac Cardiovasc Surg. 2010; 140(3): e40–e41. doi: 10.1016/j.jtcvs.2009.07.038.

8. Undas A, Kaczmarek P, Sładek K, Stępień E, Skucha W, Rzeszutko M, et al. Fibrin clot properties are altered in patients with chronic obstructive pulmonary disease: beneficial effects of simvastatin treatment. Thromb Haemost. 2009; 102(6): 1176–1182.

9. Fox CH, Johnson FB, Whiting J, Roller PP. Formaldehyde fixation. J Histochem Cytochem. 1985; 33: 845–853.

10. Stępień EŁ, Kamińska A, Surman M, Karbowska D, Wróbel A, Przybyło M. Fourier-transform infrared spectroscopy to show alterations in molecular composition of EV subpopulations from melanoma cell lines in different malignancy. Biochem Biophys Rep. 2021; 25: 100888. doi: 10.1016/j.bbrep.2020.100888.

11. Baker MJ, Hussain SR, Lovergne L, et al. Using Fourier transform IR spectroscopy to analyze biological materials. Nat Protoc. 2014; 9: 1771–1791.

12. Lee J, Hwang HS, Kim SE, et al. Impact of fixation on protein secondary structure in FTIR spectra. FASEB J. 2017; 31: 2817–2827.

13. Brandenburg E, Wulf K, Heilmann A, et al. Fixative-induced artefacts in molecular imaging and proteomic studies of human tissue. J Proteome Res. 2020; 19(7): 2857–2868.

14. Oleszko A, Dzwonek A, Wesołowski A, Dąbrowska-Szubartowicz M, Dulęba A, Chwiej J, et al. Effects of formalin fixation on tissue infrared spectra. Biomed Res Int. 2015; 2015: 245607.

15. Marzec ME, Rząca C, Moskal P, Stępień EŁ. Study of the influence of hyperglycemia on the abundance of amino acids, fatty acids, and selected lipids in extracellular vesicles using TOF-SIMS. Biochem Biophys Res Commun. 2022 Sep 24;622:30–36. doi: 10.1016/j.bbrc.2022.07.020.

16. Skalska ME, Durak-Kozica M, Stępień EŁ. ToF-SIMS revealing sphingolipids composition in extracellular vesicles and paternal β-cells after persistent hyperglycemia. Talanta. 2026;297(Pt A):128582. doi: 10.1016/j.talanta.2025.128582.

17. Skalska M. Lipid analysis in biological structures – Is time-of-flight secondary ion mass spectrometry a valuable tool in nano-lipidomics?. Bio-Algorithms and Med-Systems. (2025);21(Special Issue (OMICS innovations)):8–19. 10.5604/01.3001.0055.4327.

18. Stępniewski J, Jonas K, Magoń W, Karnaś M, Chaba W, Kopeć G. Bloodless, lytic-free, large-bore mechanical thrombectomy for the treatment of acute pulmonary embolism: the first Polish experience with the FlowTriever system. Pol Arch Intern Med. 2025; 135(2): 16880.

19. Stępniewski J, Magoń W, Podolec P, Kopeć G. The PENUMBRA Lightning 12 system for treatment of acute intermediate-high pulmonary embolism: Initial experience in Pulmonary Circulation Center Krakow, Poland. Postepy Kardiol Interwencyjnej. 2022; 18(3): 314–316.

20. Barth A, Zscherp C. What vibrations tell us about proteins. Q Rev Biophys. 2002; 35(4): 369–430.

21. Gijbels MJ, de Winther MPJ, Heeringa P, Lovric S, Wessels B, Haraldsson B. Age and lipid metabolism as determinants of atherosclerosis and arterial thrombus composition. Arterioscler Thromb Vasc Biol. 2019; 39(8): 1549–1557.

22. Jiang Y, Jin M, Wang Q, et al. Effect of aging on lipid metabolism in humans. Aging Dis. 2021; 12(1): 48–62.

23. Barth A, Zscherp C. What vibrations tell us about proteins. Q Rev Biophys. 2002; 35(4): 369–430.

24. Fagot T, Tamagne M, Le Goff W, et al. Platelet lipidomics: modern lipid profiling strategies and their relevance to atherothrombosis. J Thromb Haemost. 2019; 17(3): 365–378.

25. Cushman M, Psaty BM, Macy E, et al. Serum lipid levels and the risk of venous thrombosis. Arterioscler Thromb Vasc Biol. 2004; 24(10): 1971–1976.

26. Morelli VM, Lijfering WM, Bos MHA, et al. Lipid levels and risk of venous thrombosis. Eur J Epidemiol. 2017; 32(6): 501–508.

27. Huang Y, Liu S, Zhu Y, et al. Association between blood lipid levels and lower extremity deep venous thrombosis: a case–control study and meta-analysis. Clin Appl Thromb Hemost. 2022; 28: 10760296221121282.

28. Blat A, Dybas J, Chrabaszcz K, Bulat K, Jasztal A, Kaczmarska M, Pulyk R, Popiela T, Slowik A, Malek K, Adamski MG, Marzec KM. FTIR, Raman and AFM characterization of the clinically valid biochemical parameters of the thrombi in acute ischemic stroke. Sci Rep. 2019 Oct 29;9(1):15475. doi: 10.1038/s41598-019-51932-0.

